# Recent selection is a major force driving cancer evolution

**DOI:** 10.1101/2021.12.27.474305

**Authors:** Langyu Gu, Guofen Yang

## Abstract

Cancer is one of the most threatening diseases to humans. Understanding the evolution of cancer genes is helpful for therapy management. However, systematic investigation of the evolution of cancer driver genes is sparse. Using comparative genomic analysis, population genetics analysis and computational molecular evolutionary analysis, we detected the evolution of 568 cancer driver genes of 66 cancer types across the primate phylogeny (long timescale selection), and in modern human populations from the 1000 human genomics project (recent selection). We found that recent selection pressures, rather than long timescale selection, significantly affect the evolution of cancer driver genes in humans. Cancer driver genes related to morphological traits and local adaptation are under positive selection in different human populations. The African population showed the largest extent of divergence compared to other populations. It is worth noting that the corresponding cancer types of positively selected genes exhibited population-specific patterns, with the South Asian population possessing the least numbers of cancer types. This helps explain why the South Asian population usually has low cancer incidence rates. Population-specific patterns of cancer types whose driver genes are under positive selection also give clues to explain discrepancies of cancer incidence rates in different geographical populations, such as the high incidence rate of Wilms tumour in the African population and of Ewing’s sarcomas in the European population. Our findings are thus helpful for understanding cancer evolution and providing guidance for further precision medicine.

## Introduction

Cancer is one of the most threatening diseases with a high motality rate in humans. Somatic mutations of cancer genes can improve the fitness of tumor cells by promoting their survival, growth, and proliferation and are often under positive selection at the cellular level (Martincorena et al. 2017). Unlike somatic mutations, germline mutations of cancer genes at the individual level are usually unwelcome since mutations can be deleterious to the organism and thus affect the fitness (Fischer et al. 2011; Shendure and Akey 2015). With this logic, germline mutations of cancer genes should be dominant by purifying selection. Surprisingly, studies continuously found that cancer genes are under positive selection by comparative genomics analysis (Kang and Michalak 2015; Vicens and Posada 2018), indicating their advantageous roles during evolution. Several hypotheses have been proposed to explain the evolutionary trade-off between positive selection and cancer risks, such as sexual selection, pathogen-host interactions, and genomic compensation (Nielsen et al. 2005; Crespi and Summers 2006; Vicens and Posada 2018). However, few studies detected positive selection considering variation both among sites and branches across phylogeny. The extent to which positive selection is specific to the human lineage remains unknown.

Positive selection detection across phylogeny usually detects selection on a large time scale, i.e., usually million years (long timescale selection) (Schaschl and Wallner 2020). In contrast, detection at the population level can reflect recent selection in modern human populations. Neolithic demographic transition and the migration out of Africa have led modern humans to experience significant changes in lifestyles and living environments over the last 100,000 years (Stewart and Stringer 2012; Scerri et al. 2019). Diversified living environments and food resources accompanied by new infectious diseases brought new seletive pressures to modern human populations (Benton et al. 2021). How humans interact and adapt to changing environments continues to attract attention. Thus, the last 100,000 years are one of the most interesting time scales in human history (Sabeti et al. 2002; Sabeti et al. 2007; Schaschl and Wallner 2020). Interestingly, many human disease-related genes have been reported under positive selection due to their pleiotropic adaptive functions, such as BRCA1, SPANX, and oncogenetic viruses (Burrows et al. 2004; Kouprina et al. 2004; Lou et al. 2014). Whether genes that cause cancer, the common disease in modern humans, also show a similar signature remains unkonwn.

Although the importance of the evolution of cancer genes began to attract attention, systematic investigation of these topics in a broader view is sparse. Existing studies have mainly focused on a few cancer genes of specific cancer types or limited species across phylogeny (Huttley et al. 2000; Wildman et al. 2003; Nielsen et al. 2005; Lou et al. 2014). Besides, studies usually did not differentiate cancer driver genes and passenger genes, which could be under different selection pressures(Martincorena et al. 2017). An in-depth comparative analyses of cancer genes of different cancer types across phylogeny will give us a comprehensive picture of cancer gene evolution.

To obtain a deep understanding of the evolutionary of cancer, we detected the evolution of 568 cancer driver genes of 66 cancer types across the primate phylogeny (long timescale selection), as well as in modern human populations retrieved from the 1000 human genomics project (recent selection). We aim to answer the following question: does long timescale selection or recent selection play an important role in the evolution of cancer driver genes in humans? Our findings will have profound implications for understanding the evolution of cancer driver genes and provide clues for further precision medicine.

## Results

### Postive selection detection across the primate phylogeny

Transcript IDs of 568 cancer driver genes across 66 cancer types were retrieved from the literature published in Nature Reviews Cancer recently (Martínez-Jiménez et al. 2020). To understand the evolutionary dynamics of 568 cancer driver genes across the primate phylogeny, we obtained orthologous sequences in 24 primates using Oma inference. We further detected positive selection in different lineages under the branch-site model using PAML. Branches showing unusually large *ω* values (*ω*>100) could be due to positive selection but could also be because sequences are too divergent so that dS is saturated. We thus removed 92 genes with foreground branches showing unusually large *ω* values for downstream PAML analysis. We obtained two genes that are detected under positive selection in the human lineage. ZNF626 (ENST00000601440) belongs to Zinc finger protein coding genes with significant roles in regulatory evolution and local adaptation (Perdomo-Sabogal and Nowick 2019; Jovanovic et al. 2021). The RGPD3 gene (ENST00000409886) is related to human skull shape and face morphology determination (Wu et al. 2019). These two genes are both involved in limited cancer types, i.e., The RGPD3 gene is involved in cutaneous melanoma of the skin (CM) and the thyroid adenocarcinoma (THCA), and ZNF626 gene is involved in the nasopharyngeal cancer NSPH (Table 1).

**Table 1.** Genes under positive selection in the human lineage

| Ensembl | gene name | main function | cancer type | branch-site model |  |  |  |
| --- | --- | --- | --- | --- | --- | --- | --- |
|  |  |  |  | model A | model B | 2ln(A-B) | *W* |
| ENST00000601440 | ZNF626 | regulatory evolution | NSPH | -6898.8 | -6906.9 | 16.32 | 8.67 |
| ENST00000409886 | RGPD3 | skull shape and face morphology | CM, THCA | -12854 | -12881 | 53.66 | 70.6 |

In contrast, many genes are under positive selection in ancestral lineages across the primate phylogeny (Fig. 1, Supplementary File 1).

**Figure 1.**
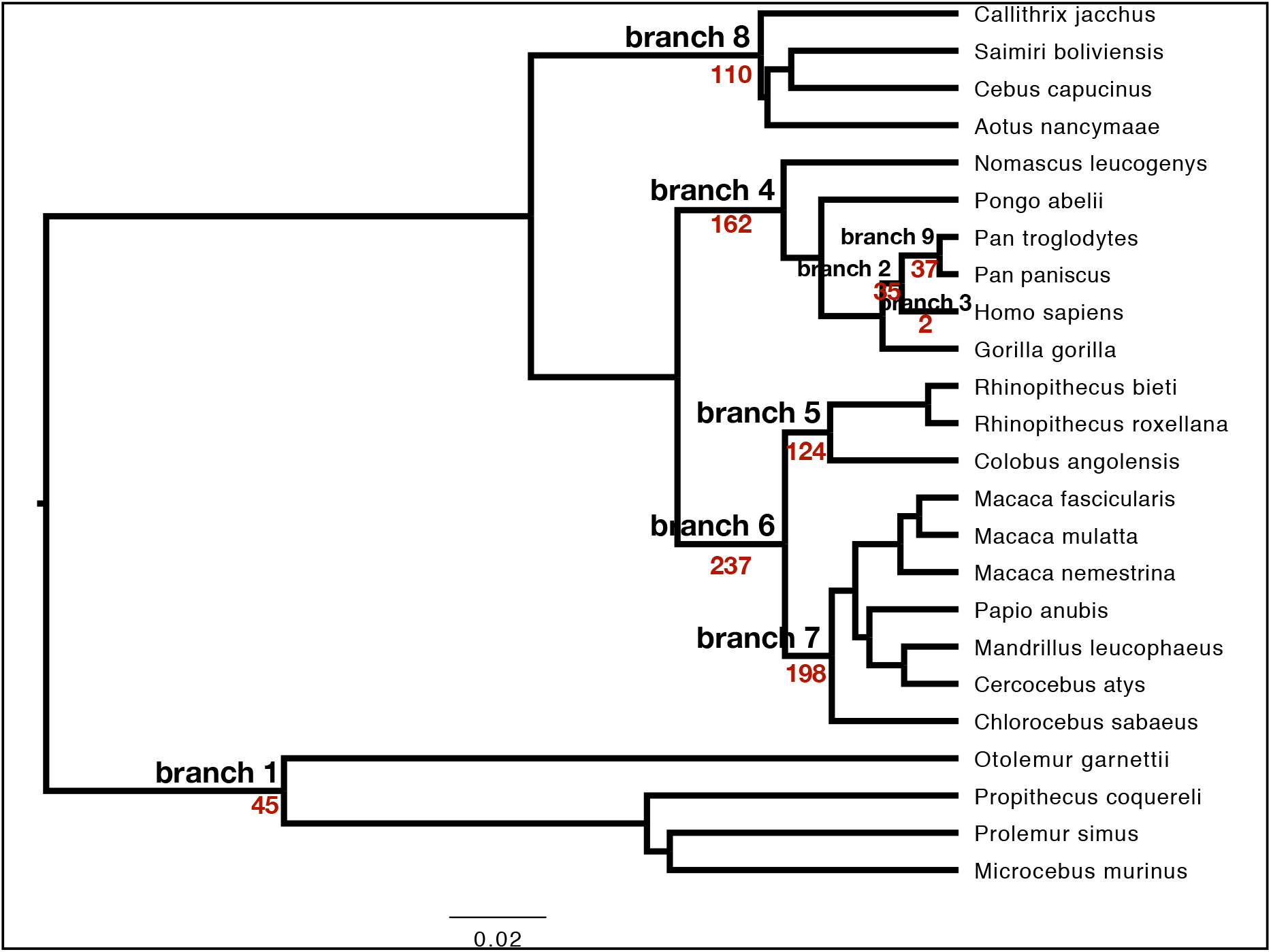
Genes under positive selection using a branch-site model in PAML across the primate phylogeny. Target foreground branches were labeled. The number of genes under positive selection was given in the target foreground branch in red. The species tree was retrieved from the literature (Upham et al. 2019). The unrooted tree was used for PAML analysis.

### Positive selection detection in modern human populations

We inferred SNPs that were under positive selection based on the following criteria: 1) the absolute value of LD-based test statistics iHS >2 in the population detected. 2) SNPs with iHS>2 also showed high pairwise Fst (>0.20) between the detected population and the other population. We thus found 77 cancer driver genes under positive selection in total in different populations (Supplementary File 2).

Among these 77 genes, ten genes have been reported under positive selection, including host-pathogen interaction-related genes (PARP4(Daugherty et al. 2014; Gossmann and Ziegler 2014), SH2B3(Coenen et al. 2009; Zhernakova et al. 2010; Aaron 2011)), neuron-related genes (FAT1(Sato and Kawata 2018), NIN(Montgomery and Mundy 2012)), pigmentation-related genes (HERC2(Wilde et al. 2014; Cheng et al. 2017), EGFR(Lao et al. 2007)), metabolic-related genes (UGT2B17(Xue et al. 2008)), the well-known tumor suppressor gene TP53(Levine 2020), and the two genes mentioned above, i.e., Skull shape and facial morphology-related gene RGPD3 (Wu et al. 2019) and regulatory evolution-related ZNF gene ZNF626(Perdomo-Sabogal and Nowick 2019; Jovanovic et al. 2021). Gene enrichment analysis revealed that these 77 positively selected genes were involved in immune-related pathways, p53 pathways, neuron-related pathways, angiogenesis, the Insulin/IGF pathway, the PDGF pathway, the FGF pathway, the VEGF pathway, the EGF pathway, and the Wnt signaling pathway (Fig.2).

**Figure 2.**
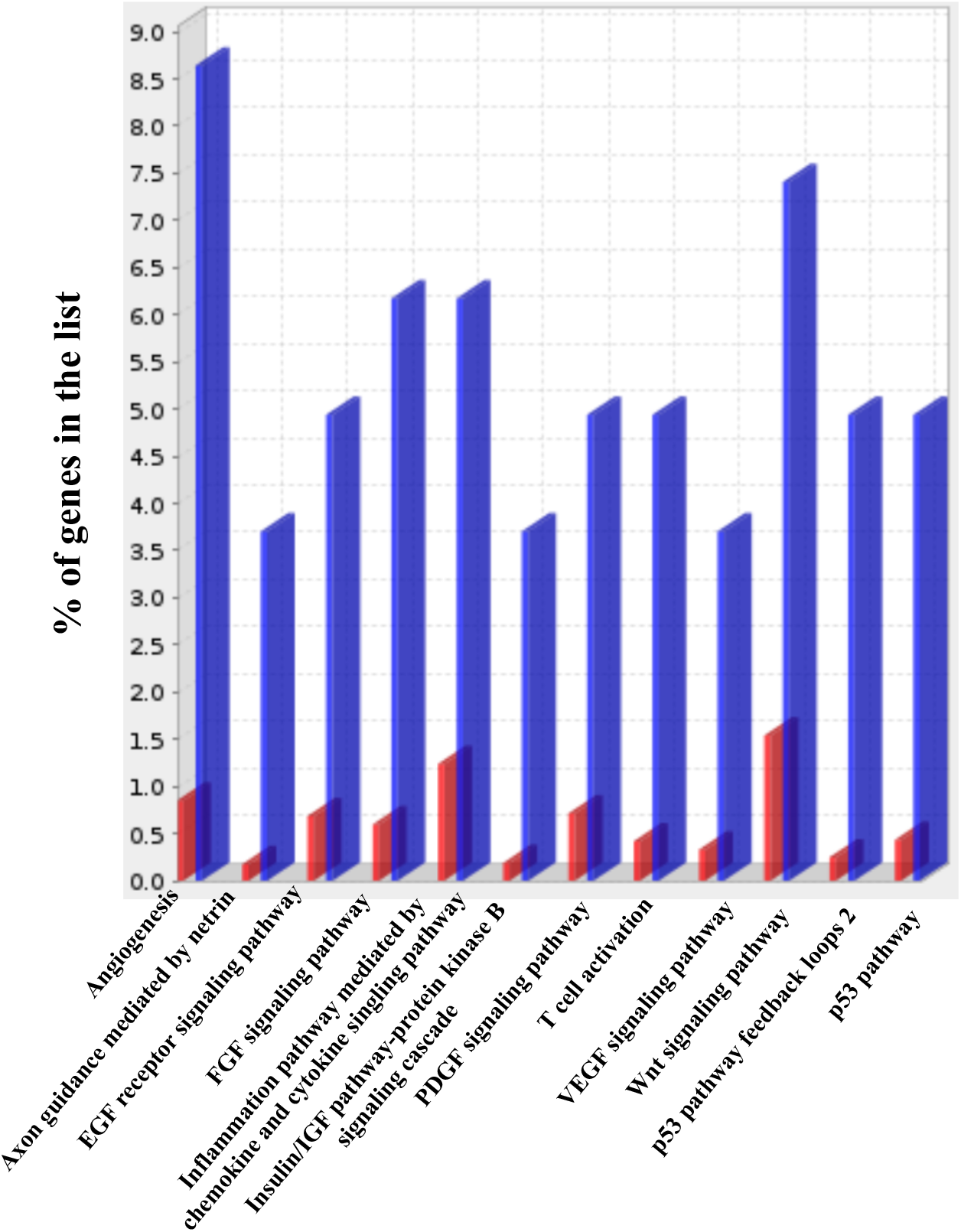
Pathway enrichment of positively selected cancer driver genes in modern human populations. % of gene list in the category is calculated for each testing list as: genes for the category/total genes in the list * 100. The red bar represents the reference human genome. The blue bar represents the target positively selected genes. It displays only results for FDR *p*<0.05.

Cancer driver genes are under positive selection in different populations (Fig. 3, Supplementary File 2). There are 27 genes corresponding to 39 cancer types under positive selection in the African population, 26 genes corresponding to 47 cancer types under positive selection in the East Asian population, 35 genes corresponding to 61 cancer types under positive selection in the European population, and 6 genes corresponding to 11 cancer types under positive selection in the South Asian population. Note that the African population shows the largest extent of divergence with other populations. On the one hand, among 27 positively selected genes in the African population, there are 15 genes showing high Fst compared with the European population, 24 genes showing high Fst compared with the East Asian population, 17 genes showing high Fst compared with the South Asian population. On the other hand, other populations showed large numbers of positively selected genes with high Fst compared with the African population. For example, there are 32 out of 35 positively selected genes in the European population showing high Fst compared with the African population; 26 out of 26 positively selected genes in the East Asian population showing high Fst compared with the African population; and 6 out of 11 positively selected genes showing high Fst compared with the African population.

**Figure 3.**
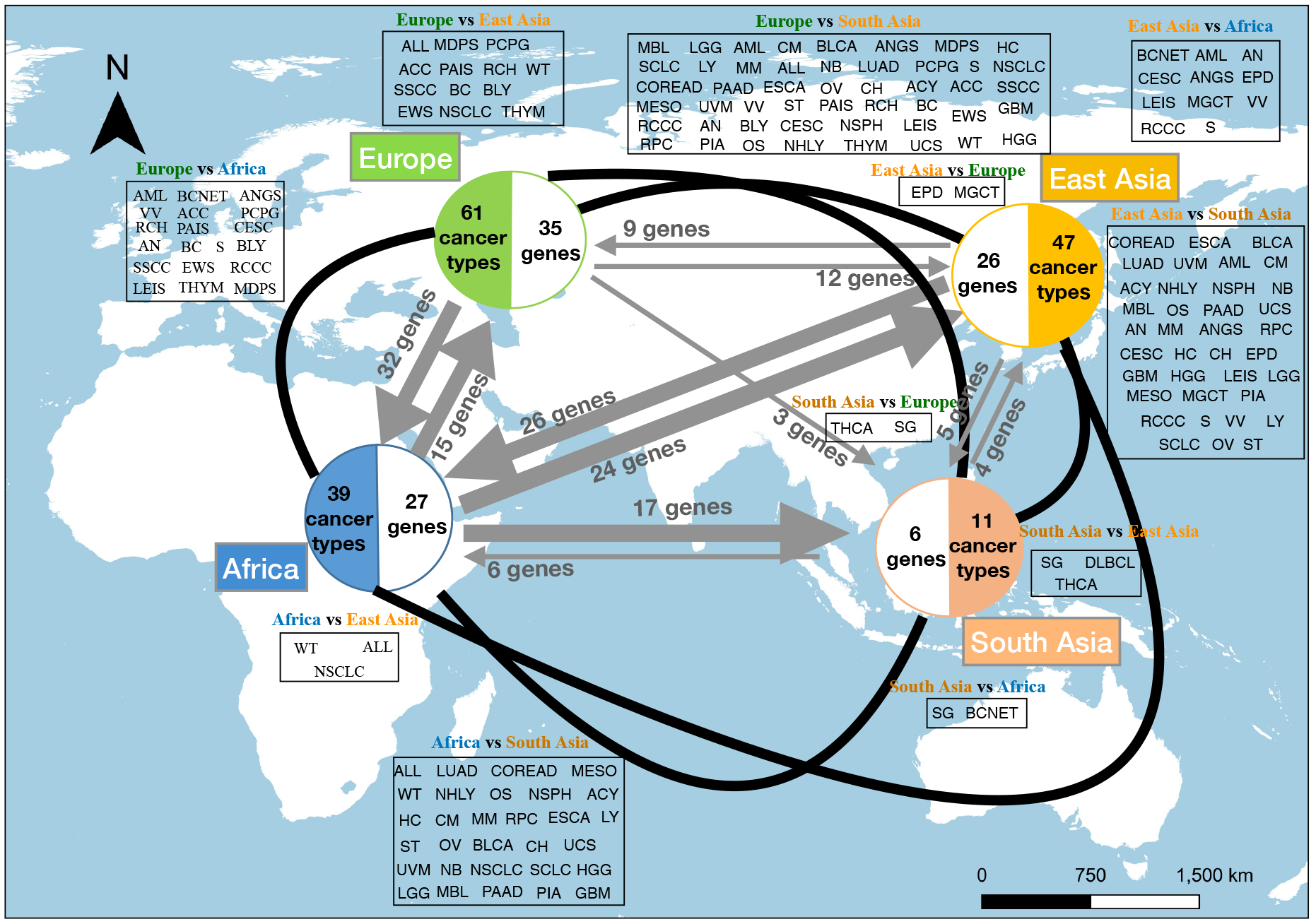
Genes under positive selection in modern human populations, i.e., the African population, the European population, the East Asian population, and the South Asian population. Arrows showed positively selected genes with iHS>2 in the local population and Fst >0.20 between populations. There are 27 genes corresponding to 39 cancer types under positive selection in the African population, 26 genes corresponding to 47 cancer types under positive selection in the East Asian population, 35 genes corresponding to 61 cancer types under positive selection in the European population, and 6 genes corresponding to 11 cancer types under positive selection in the South Asian population. The African population showed the largest extent of divergence with other populations. The European population has the largest numbers of cancer types (61 cancer types) whose driver genes are under positive selection, followed by East Asian (47 cancer types) and African (39 cancer types) populations. The South Asian population has the lowest numbers of cancer types (11 cancer types). The corresponding cancer types whose driver genes are under positive selection showing population-specific patterns are given in boxes. Abbreviations of cancer types can be found in the Supplementary File 3.

When took account the corresponding cancer types of positively selected cancer driver genes in each population, we found that the European population has the largest numbers of cancer types (61 cancer types), followed by East Asian (47 cancer types) and African (39 cancer types) populations. The South Asian population has the lowest numbers of cancer types (11 cancer types). These cancer types also exhibited population-specific patterns. For example, driver genes of cancer types of WT (Wilms tumor), ALL (acute lymphoblastic leukaemia) and NSCLC (non small cell lung cancer) are under positive selection in the African popualtion but not in the East Asian population. Driver genes of cancer types of SG (salivary gland cancer) and BCNET (small intestine cancer neuroendocrine) are under positive selection in the South Asian population but not in the African population. Driver genes of cancer types of SG, DLBCL (diffuse large B cell lymphoma) and THCA (thyroid adenocarcinoma) are under positive selection in the South Asian population but not in the East Aisan population. More details can be found in Fig. 3. Abbreviations of cancer types can be found in the Supplementary File 3.

## Discussion

Using comparative genomic analysis, population genetic analysis, and computational molecular evolutionary analysis across the primate phylogeny and in modern human populations, we found that recent selection, rather than long timescale selection, has significant effects on the evolution of cancer driver genes in humans. Cancer driver genes related to morphological traits and local adaptation are under positive selection in different human populations. The corresponding cancer types of these positively selected genes showed population-specific patterns, providing clues to explain cancer incidence rates discrepancy in different human populations. These findings are helpful for understanding cancer evolution and providing clues for further precision medicine.

A few studies have already conducted a molecular evolutionary analysis of cancer genes across primates and found many cancer genes are under positive selection in primates (Lou et al. 2014). We also found tens of hundreds of genes under positive selection in primates (Fig. 1). This indicates that these genes indeed play advantageous roles during the early evolution of primates. The genetic background of many cancer genes can be traced back to before human lineage itself. For example, some cancer genes are essential biological genes controlling basic biology, such as DNA damage repair, cell division (O’Connell 2010; Wu et al. 2020; Zahir et al. 2020), or genes associated with adaptation (Muller 2017; Shi et al. 2019). As we mentioned in the introduction section, several hypotheses have already been proposed to explain this phenomenon.

However, when we focused on positive selection in the human lineage under a branch-site model, we found only two cancer driver genes under positive selection (Fig. 1). The reason could be that their harmful effects on human evolution outweighed their advantageous roles in the early evolution of primates. Indeed, massive changes occurred during the evolution of human beings, such as upright walking (Niemitz 2010; Gruss and Schmitt 2015; Zirkle and Lovejoy 2019) and brain development (Kanton et al. 2019), which can induce different selection pressures in the evolution of the human lineage. In addition, nonhuman primates might have other cancer genetic bases or anticancer mechanisms, which are not rare in mammals, including mole rats, elephants and whales (Seluanov et al. 2018). Comparisons among species provide a snapshot of selected events that have been unfolding over long timescales. The few numbers of cancer driver genes under positive selection in the human lineage suggests that long timescale positive selection plays a minor role in driving the evolution of human cancer.

In contrast, 77 cancer driver genes are under recent positive selection in modern human populations. Most of these genes are related to host-pathogen interactions, neuron development, pigmentation and metabolism, which are involved in human-environmental interactions(Lao et al. 2007; Xue et al. 2008; Coenen et al. 2009; Zhernakova et al. 2010; Aaron 2011; Montgomery and Mundy 2012; Daugherty et al. 2014; Gossmann and Ziegler 2014; Wilde et al. 2014; Cheng et al. 2017; Sato and Kawata 2018; Levine 2020). Many genes showed population-specific positive selection patterns (Fig. 3), suggesting their adaptations to different environmental conditions. Significant effects of condition changes in modern human populations instead of long timescales evolution are also found in the evolution of testis-related genes (Schaschl and Wallner 2020). Food consumptions, environmental pollution, living stress and culture affecting diseases in modern human populations have increasingly attracted attention(review in (Benton et al. 2021)). The numbers of cancer driver genes under positive selection show geographical differences among populations, with the African population showing the largest extent of divergence with other populations. These suggest that cancer driver genes are involved in local adaptation and could be related to the migration *out of Africa* during modern human evolution (Stewart and Stringer 2012; Scerri et al. 2019).

It is worth noting that the corresponding cancer types whose driver genes are under positive selection show population-specific patterns (Fig. 3). Wilms tumor (WT) is more common in Africa than in East Asia (Leslie et al. 2021), which is in consitent with our findings that driver genes of WT are under positive selection in the African population but not in the East Asian population (Fig. 3). Another case is Ewing’s sarcomas (EWS), which occurs more popular in the European population than in African or Asian populations. Genetic component is believed to be a key factor for developing this cancer (Worch et al. 2010). Our study gives a support of this opinion, giving that driver genes of EWS are found under positive selection in the European population, but not in African and Asian populations (Fig. 3). Noticeably, the South Asian population has the lowest numbers of cancer types whose driver genes are under positive selection (11 cancer types). This gives a good explaination why the South Asian population usually has low cancer incidence rate (Tran et al. 2018). Our findings that varied cancer driver genes are under different selection pressures in different populations can thus help explain geographical disparities of cancer incidence rates, although other factors such as environmental conditions and living styles are also important (Bahnassy et al. 2020). These results again, suggest that recent positive selection is a major force driving the evolution of cancer in humans.

A genetic disease can occur as a byproduct of evolution. For instance, the seemingly human-specific disease of schizophrenia and the greater human susceptibility to Alzheimer’s disease may be a byproduct of the human specialization for higher cognitive function (Vamathevan et al. 2008). This can also be applied to cancer to humans. Cancer driver genes under recent positive selection in human populations we found here suggest that pleiotropic effects should be under consideration during precision cancer medicine.

## Materials and Methods

### Orthologous sequences inference

Transcript IDs of 568 cancer driver genes across 66 cancer types were retrieved from the literature published in Nature Reviews Cancer recently (Martínez-Jiménez et al. 2020). The corresponding sequences retrieved from Ensembl Biomart were used as queries in a tblastx search with *e* −0.001 against each species’s related genome-wide coding seqeunces (cds). Cds of the first top tblastx hit of each queried transcript were extracted for the corresponding species. Oma was used to perform further orthologous sequence inference with default parameters. A species tree was retrieved from the literature (Upham et al. 2019), and *Mus musculus* was used as the outgroup.

### Positive selection detection across the phylogeny of primates

The rates *ω* of nonsynonymous (dN) to synonymous (dS) substitutions can vary over sites and time. Positive selection detection was based on the rates *ω* calculated using a branch-site model with codeml in PAML. The branch-site model considers the variation in *ω* among sites and acorss branches to detect positive selection affecting sites along the target lineage (foreground branch). If a nonsynonymous mutation is less likely to be tolerated during evolution, *ω* will be <<1, meaning that purifying selection takes effects. If there is no selection pressure (neutral selection), *ω* will be equal to 1. If changes of nonsynonymous mutations have beneficial effects and are favored by selection, *ω* will be >>1, indicating that they are under positive selection.

Multiple sequence alignment was conducted using MAFFT within T-coffee. An unrooted species tree retrieved from the literature (Upham et al. 2019) was used for the calculation. Nine branches were set as foreground branches to run the branch-site model separately (Fig. 1). Bonferroni correction was used for multiple test corrections. Parameter settings and statistical significance tests were the same as in the literature (Gu and Xia 2019).

### Positive selection detection in modern human populations

The integrated haplotype score method (iHS) was applied to identify recently positively selected genes in different human populations. The principle of this method is that under the condition of neutral evolution, the high-frequency rise rate of the selected allele is faster than expected. The long-range association with the surrounding loci will not have enough time to be eliminated by recombination. Therefore, by measuring the association attenuation between core haplotypes and alleles at different distances, i.e., extended haplotype homozygosity (EHH), we can detect signals of recent positive selection in favor of variants that have not yet reached fixation. One positive selection feature is finding a core haplotype combining high frequency and high EHH compared with other core haplotypes at the site. Therefore, various core haplotypes at one locus can be used as internal controls to adjust the heterogeneity of the local recombination rate.

Phased genetic data of the 1000 genomes project of modern human populations were retrieved from ftp.1000genomes.ebi.ac.uk/vol1/ftp/release/20130502/. We analyzed larger georgraphical areas, including Africa, East Aisa, Europe, and South Asia. Details can be found in the Supplementary File 4. PLINK and VCFtools were used to process variant call format (VCF) files for all chromosomes. Only biallelic SNPs with minor allele frequency (MAF)>0.05 and indels removed were considered in this study. The human genome GTF file from Ensembl was used to retrieve genomic location of cancer genes. The integrated haplotype score test (iHS) implemented in selscan was used to detect genome-wide positive selection with default parameters. The unstandardized iHS scores were normalized in frequency bins across the entire genome using the script *norm* implemented in the selscan program. Pairwise Fst was calculated for each SNP using the Weir & Cockerham Fst calculation implemented in VCFtools.

### Pathway enrichment analysis

The pathway enrichment analysis of target positively selected cancer driver genes was conducted using the Gene Ontology Resource (http://geneontology.org/) with the human genome as the background. Fisher’s exact test was conducted, and false discovery rates (<0.05) were calculated to enrich significant pathways in the Panther pathways database.

## Acknowledgements

We thank Dr. Juan Felipe Ortiz, Dr. Haosen Li, Dr. Cheng Huang and Prof. Canwei Xia for the support of data analyses. This work was supported by the Fundamental Research Funds for the Central Universities (20lgpy109) to LG and the National Natural Science Foundation of China (81772769) to GY; Data analyses was supported by National Supercomputer Center in Guangzhou, China. The English language was edited by Elsevier Language Editing Services.

## Author Contributions

LG and GY designed the study. LG did the analysis. LG and GY wrote the manuscript.

## Data Availability Statements

Detail software and corresponding parameters were given in the Methods.

## Supplementary Files

Supplementary File 1 Genes under positive selection under branch-site model with PAML across the primate phylogeny.

Supplementary File 2 Recent positive selection detection within human populations and Fst comparisons between human populations.

Supplementary File 3 Abbreviations of 66 cancer types.

Supplementary File 4 IDs of population data retrieved from the 1000 human genomes project.

